# Role of substrate recognition in modulating strigolactone receptor selectivity in witchweed

**DOI:** 10.1101/2020.07.28.225722

**Authors:** Jiming Chen, Alexandra White, David C. Nelson, Diwakar Shukla

## Abstract

Witchweed, or Striga *hermonthica*, is a parasitic weed that destroys billions of dollars worth of crops globally every year. Its germination is stimulated by strigolactones exuded by its host plants. Despite high sequence, structure, and ligand binding site conservation across different plant species, one strigolactone receptor in witchweed (*Sh* HTL7) uniquely exhibits a picomolar EC50 for downstream signaling. Previous biochemical and structural analyses have hypothesized that this unique ligand sensitivity can be attributed to a large binding pocket volume in *Sh* HTL7 resulting in enhanced ability to bind substrates. Additional structural details of the substrate binding process can help explain its role in modulating the ligand selectivity. Using long-timescale molecular dynamics simulations, we demonstrate that mutations at the entrance of the binding pocket facilitate a more direct ligand binding pathway to *Sh* HTL7, whereas hydrophobicity at the binding pocket entrance results in a stable “anchored” state. We also demonstrate that several residues on the D-loop of *At* D14 stabilize catalytically inactive conformations. Finally, we show that strigolactone selectivity is not modulated by binding pocket volume. Our results indicate that while ligand binding is not the sole modulator of strigolactone receptor selectivity, it is a significant contributing factor. These results can be used to inform the design of selective antagonists for strigolactone receptors in witchweed.

## 1 Introduction

Strigolactones are a class of plant hormones responsible for regulating shoot branching and root architecture in plants^1–4^. They have also been found to induce seed germination in the parasitic *Striga* genus^5^. Estimates of global crop losses due to *Striga* parasites are in excess of $10 billion per year, warranting a need for effective *Striga* control^6^. Strigolactone perception is controlled by a family of proteins called DWARF14, which possess a conserved *α*-*β* hydrolase fold with a hydrophobic cavity in which the substrate binds. D14 and its closely related homolog KAI2, contain a strictly conserved Ser-His-Asp catalytic triad (Fig. 1a). Strigolactone signalling responses are believed to be dependent on enzymatic hydrolysis of the substrate and subsequent covalent modification of the enzyme by a hydrolysis product^7–10^. Following hydrolysis, the receptor undergoes a large conformational change that enables it to associate with MAX2 and SMXL proteins, which are then ubiquitinated and degraded by the proteasome^11^. Recent evidence has also suggested that signal can be transduced by intact strigolactone molecules^12^, and that MAX2 proteins may act as a repressor of strigolactone hydrolysis^13^.

**Fig. 1.**
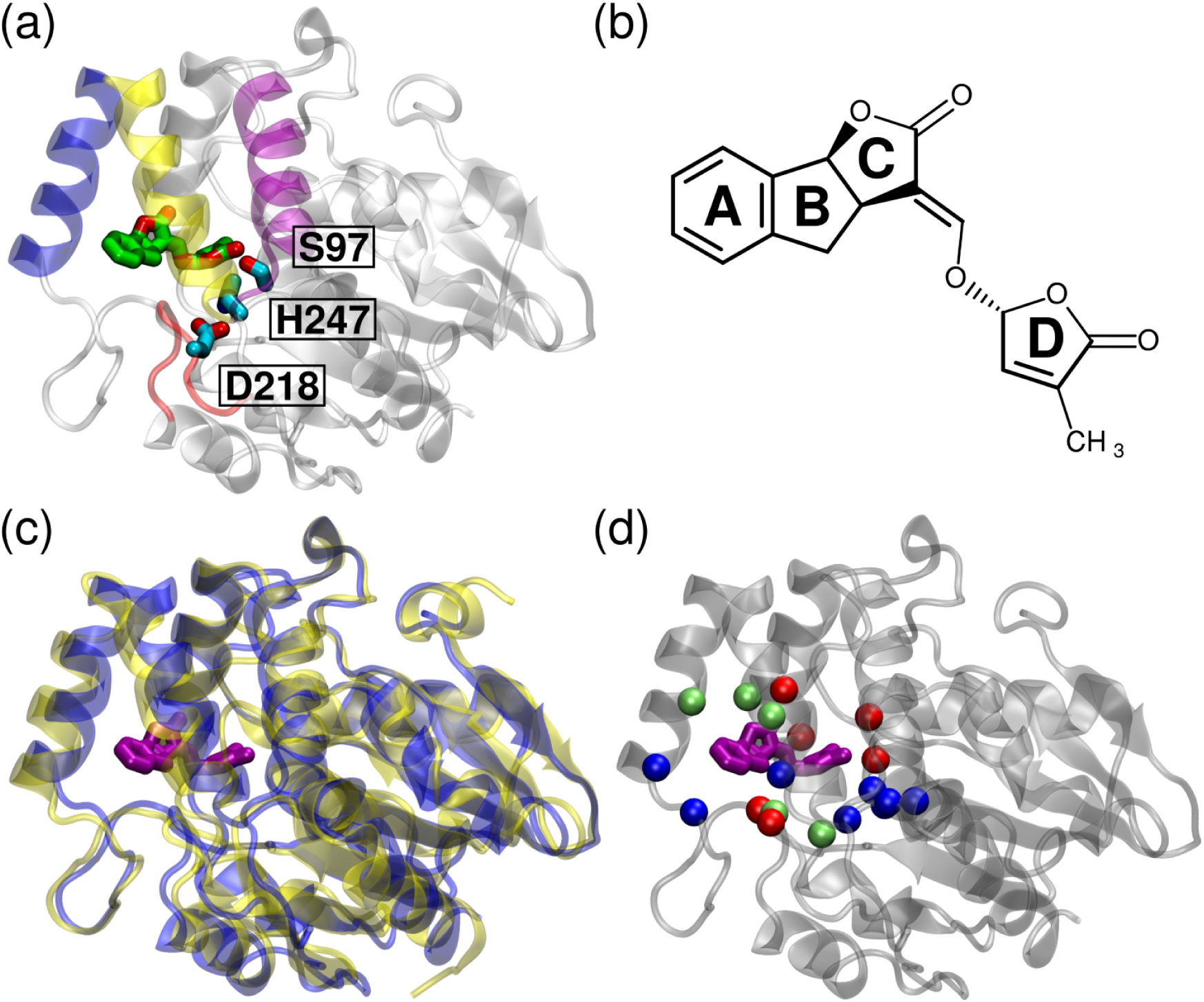
(a) Structure of *Arabidopsis thaliana* strigolactone receptor *At* D14 in complex with strigolactone analog GR24 (green). The T1, T2, and T3 helices are shown in blue, yellow, and purple, respectively, and the D-loop is shown in red. The serine-histidine-aspartate catalytic triad is shown in cyan. (b) Structure of synthetic strigolactone analog GR24^5DS^, which possesses the sterochemistry of naturally occuring strigolactones. (c) Structural alignment of *At* D14 (blue) and *Sh*HTL7 (yellow). Bound GR24 is shown in purple. (d) Similarity of binding pocket residues. Blue spheres indicate conserved residues between *At* D14 and *Sh*HTL7, red spheres indicate similar residues, and green spheres indicate different residues. Full sequence and secondary structure alignments are shown in Fig. S1.

Despite 44% sequence identity between *Arabidopsis thaliana* and *Striga hermonthica* strigolactone receptors, 78% sequence similarity, and a highly conserved structure between different species (Fig. 1c), one receptor in *Striga hermonthica, Sh*HTL7, uniquely exhibits a picomolar range EC50 for downstream signaling for inducing a germination response, compared to micromolar ranges for other strigolactone receptors^14^. An evolutionary analysis by Conn *et al*. revealed that *Sh*HTL proteins evolved the ability to perceive strigolactone via convergent evolution^15^. *Sh*HTL proteins are paralogs of KAI2 proteins, which perceive seed germination stimulants in plants and evolved strigolactone sensitivity independently of D14 proteins^15^. KAI2 proteins are generally grouped into three clades, the KAI2c (conserved) clade which is the most KAI2-like and has sensitivity to karrikins but not strigolactones, the KAI2i intermediate clade, and the divergent KAI2d clade that is strigolactone-sensitive but not karrikin-sensitive. Further studies have hypothesized that the high strigolactone sensitivity found in several members of the divergent clade of *Sh*HTL proteins, which includes *Sh*HTL7, can be attributed to their larger binding pocket volume compared to other members of the D14/KAI2 superfamily of proteins^14, 16^. Additionally, an isothermal titration calorimetry and crystallography study by Burger et. al suggested that the T2-T3 loop of KAI2 proteins is able to modulate pocket size which in turn is able to influence binding affinity^17^. However, this hypothesis relies on pocket volumes computed from crystal structures, which can only provide static “snapshots” of the protein. In an aqueous environment, the pocket volume is likely to fluctuate due to conformational flexibility of the protein.

Alternatively, differences in the substrate binding mechanism can contribute to enhanced signaling ability in one protein over the other. Differences in the substrate binding process can enhance signaling in two ways: (i) A higher binding affinity for the ligand can increase the residence time of the ligand in the pocket, leading to increased probability of enzymatic hydrolysis occurring, or (ii) a lower free energy barrier of binding can enhance the rate of binding, thus enhancing the apparent rates of subsequent steps. Characterizing the role of these effects in producing the uniquely high sensitivity of *Sh*HTL7 requires a detailed structural and dynamical characterization of the binding process. While structures of the protein-ligand complexes could provide insights into differences in binding affinity, mechanistic details of the binding process can additionally determine the effects of sequence differences in residues outside the binding pocket on ligand binding. The only currently available crystal structure of a strigolactone-bound D14 protein is a structure of *Os*D14, the *Oryza sativa* ortholog of *At* D14 (*∼*74% sequence identity), bound to GR24, a synthetic strigolactone analog^18^. There is uncertainty surrounding the accuracy of this as well as other strigolactone receptor crystal structures bound to various ligands due to low electron density of ligands within the binding pocket^19^, and there is an inherent lack of dynamical information contained in crystal structures. Making direct biophysical measurements on the binding process is also particularly challenging for strigolactone receptors since it is known to hydrolyze its ligand. This coupling of binding and hydrolysis makes it difficult to elucidate the effects of substrate binding on signaling independently of subsequent steps. A powerful method that can be used to characterize the strigolactone binding process is molecular dynamics (MD) simulations^20–26^. When used with Markov state models (MSMs), simulations can provide detailed kinetic and thermodynamic information about ligand binding processes at atomic-level resolution^27–32^. Furthermore, MSMs allow us to perform analysis on a large number of short simulations rather than a single long simulation^33, 34^, which greatly decreases the time required to obtain sufficient data. MD simulations have previously been used to characterize other conformational dynamics and substrate binding in other plant proteins^20, 21, 24–26^.

Recently, Hu *et al*. employed biased MD simulations to characterize the mechanism of the smoke-derived compound KAR_1_ to *At* KAI2^26^, a homolog of *At* D14 (*∼*50% sequence identity). However, a limitation to this study is the biasing methods that were used have an inherent assumption that the ligand can only bind in a single binding pose and via a single pathway. Here, we employ long timescale (*∼*400 *µ*s aggregate) unbiased MD simulations, allowing for a high resolution, dynamical view of the substrate recognition mechanisms in *At* D14 and *Sh*HTL7. We demonstrate that *Sh*HTL7 is more efficient at binding GR24 and is also more effective at positioning GR24 for hydrolysis than *At* D14. Additionally, we show that while differences in the ligand binding process do contribute to the high ligand sensitivity in *Sh*HTL7, these differences are not caused by the difference in the crystal structure pocket volume.

## 2 Results

### Free energy profile of the binding process

Using *∼*200 *µ*s aggregate of MD simulations each on GR24 binding to *At* D14 and *Sh*HTL7, we computed the free energy landscapes of the complete ligand binding processes (Fig. 2). These landscapes were projected onto A-ring-catalytic serine distance and D-ring-catalytic serine distance for the purpose of distinguising different binding modes of the ligand. Free energy minima discernable from these landscapes are the bound state (*α*), consistent with the crystal structure of GR24-bound *Os*D14 (PDB 5DJ5)^18^; an inverse bound state (*β*), and an “anchored” intermediate state (*γ*). The most stable minimum for both proteins corresponds to the bound state with the butenolide ring of the ligand oriented into the binding pocket and close to S97/95 of the catalytic triad. Both *At* D14 and *Sh*HTL7 are also capable of binding GR24 in an inverse pose, in which the A-ring is oriented into the pocket and the butenolide ring (D-ring) is oriented toward the pocket entrance. The canonical model of strigolactone signaling involves a catalytic mechanism in which S97/95 nucleophilically attacks the ligand upon the Dring^7, 8^, indicating that this inverse-bound pose is likely catalytically inactive, and thus signaling incompetent.

**Fig. 2.**
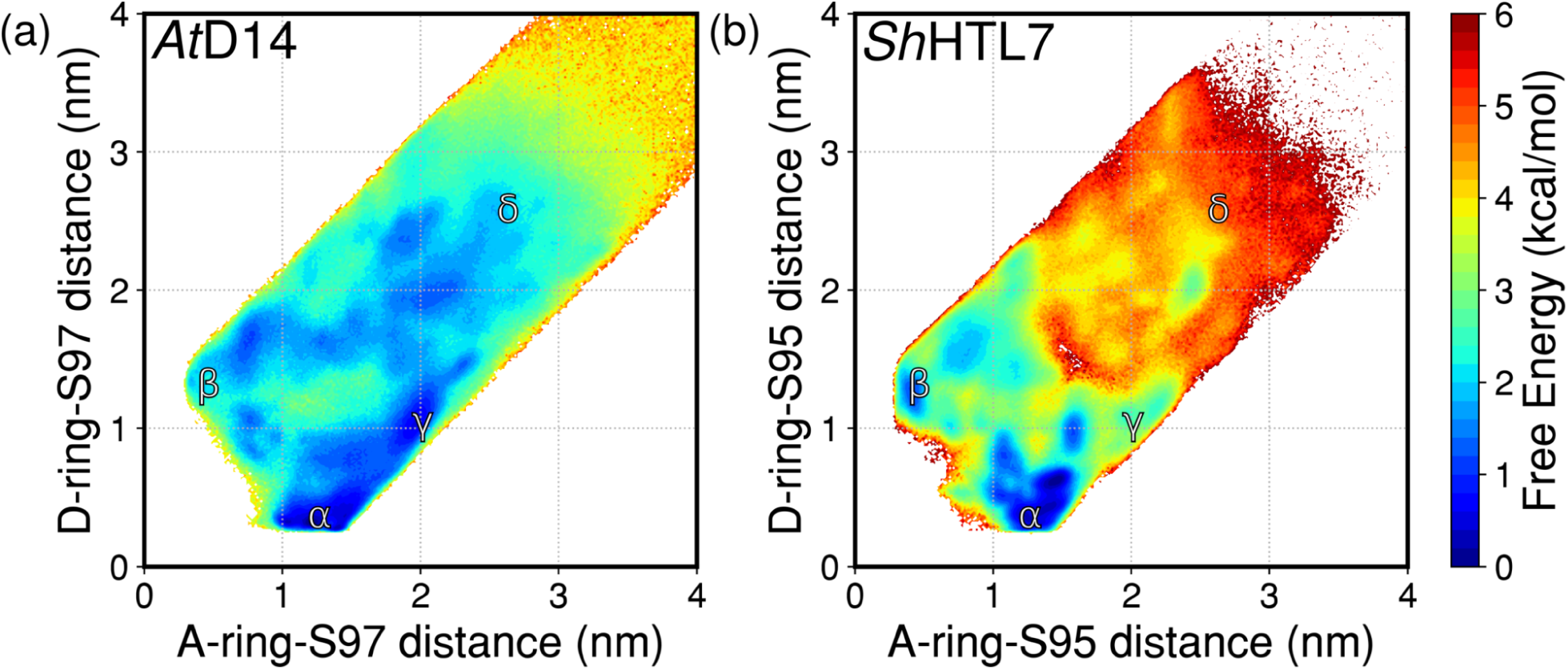
Free energy landscapes of GR24 binding to (a) *At* D14 and (b) *Sh*HTL7. Labeled states are *α*: Bound state, *β*: Inverse bound state, *γ*: Anchored state, *δ*: Unbound states

Using the method in Buch *et al*.^27^, *we calculated the free energy for GR24 binding to be -5.5 kcal/mol in At* D14 and -5.7 kcal/mol in *Sh*HTL7. Free energy landscapes with respect to the slowest motions in the binding process are shown in Fig. S2. A previously reported dissociation constant (*K*_*d*_) for GR24 binding to *At* D14 based on an isothermal titration calorimetry measurement is 0.30*±*0.02 *µ*M, which corresponds to a free energy of -8.7 kcal/mol at the experimental temperature of 293 K^35^. These free energy values were computed using the equation Δ*G* = *−RT* ln *K*_*d*_, where *T* is the temperature at experimental conditions and *K*_*d*_ is the reported dissociation constant. However, this value is likely the free energy associated with both binding and hydrolysis because a significant positive entropy change (+19.5 cal/mol*K) upon binding is reported. Ligand binding is expected to have a negative entropy change associated with loss of configurational entropy of the ligand, but a hydrolysis reaction in which GR24 is split into its ABC-ring and D-ring would more likely yield a positive entropy change. The *K*_*d*_ for GR24 binding to *Sh*HTL7 is estimated to be 0.92*±*0.01 *µ*M based on microscale thermophoresis assay and 0.39*±*0.05 *µ*M based on a tryptophan fluorescence assay, which correspond to a free energies of -8.7 kcal/mol and -8.2 kcal/mol, respectively, at 298K^36^.

### Hydrophobic to polar mutations at pocket entrance destabilize anchored intermediate state in *Sh* HTL7

A notable difference in the binding pathways is the stability of the “anchored” intermediate state (*γ*). Based on the free energy landscapes, the anchored intermediate state is *∼*1.5 to 2 kcal/mol more stable in relation to the bound minimum in *At* D14 than in *Sh*HTL7. This indicates that the ligand is more likely to interact with the pocket entrance during the binding process in *At* D14 than in *Sh*HTL7. To further investigate the pocket entrance-anchoring observed in *At* D14, we computed per-residue ligand contact probabilities for both *At* D14 and *Sh*HTL7 (Fig. 3a,b). In agreement with the free energy landscapes of the binding process, the regions of highest ligand contact probabilities in both proteins were the interior of the binding pocket. Additionally, the region directly outside the binding pocket shows considerably higher ligand contact probability in *At* D14 than in *Sh*HTL7. This further indicates the presence of a stable interaction between the ligand and a patch of T1 and T2 helix residues outside the binding pocket. A comparison of residues in contact with the ligand in the *At* D14 anchored state and their corresponding residues in *Sh*HTL7 is shown in Fig. 3c,d. Four residues on the T1 and T2 helices are mutated from hydrophobic to polar residues between *At* D14 and *Sh*HTL7: V144 (T142), A147 (S145), A154 (S152), and F159 (T157). Increased polarity at the pocket entrance prevents stable hydrophobic interactions from forming with the ABC rings of GR24. Since interactions between GR24 and the pocket entrance are largely hydrophobic, these mutations are likely to destabilize the anchored state in *Sh*HTL7, thus leading to enhanced binding kinetics. This is consistent with the observation that a G158E mutant of *At* D14 displayed increased hydrolytic activity toward GR24 despite becoming signalling inactive^7^. Additionally, in a recent study introducing a femtomolar-range suicide germination compound for *Striga*, several mutations at the binding pocket entrance resulted in an increase in IC50 for competitive binding of the compound to SPL7^37^. While this is not directly comparable since the measurements were done with a different ligand, it nonetheless supports the hypothesis that residues at the pocket entrance play an important role in ligand binding. Using the ConSurf server^38^, we also computed the site conservation of these four residues among homologs of *At* D14 and *Sh*HTL7. Most frequent residues occupying the four pocket entrance sites among *At* D14 and *Sh*HTL7 are shown in Fig. 3e and f. The four sites show high conservation among both sets of homologs. However, while the pocket entrance residues in *At* D14 all match the most frequent residues of the given sites, the pocket entrance residues in *Sh*HTL7 are all less common substitutions. Notably, the most common residues at pocket entrance sites in *Sh*HTL7 homologs are hydrophobic, as in *At* D14, indicating that polarity at the pocket entrance is not a common feature even among close homologs of *Sh*HTL7. Parameters for the ConSurf calculation can be found in Table S1, and conservation scores and residue lists can be found in Table S2.

**Fig. 3.**
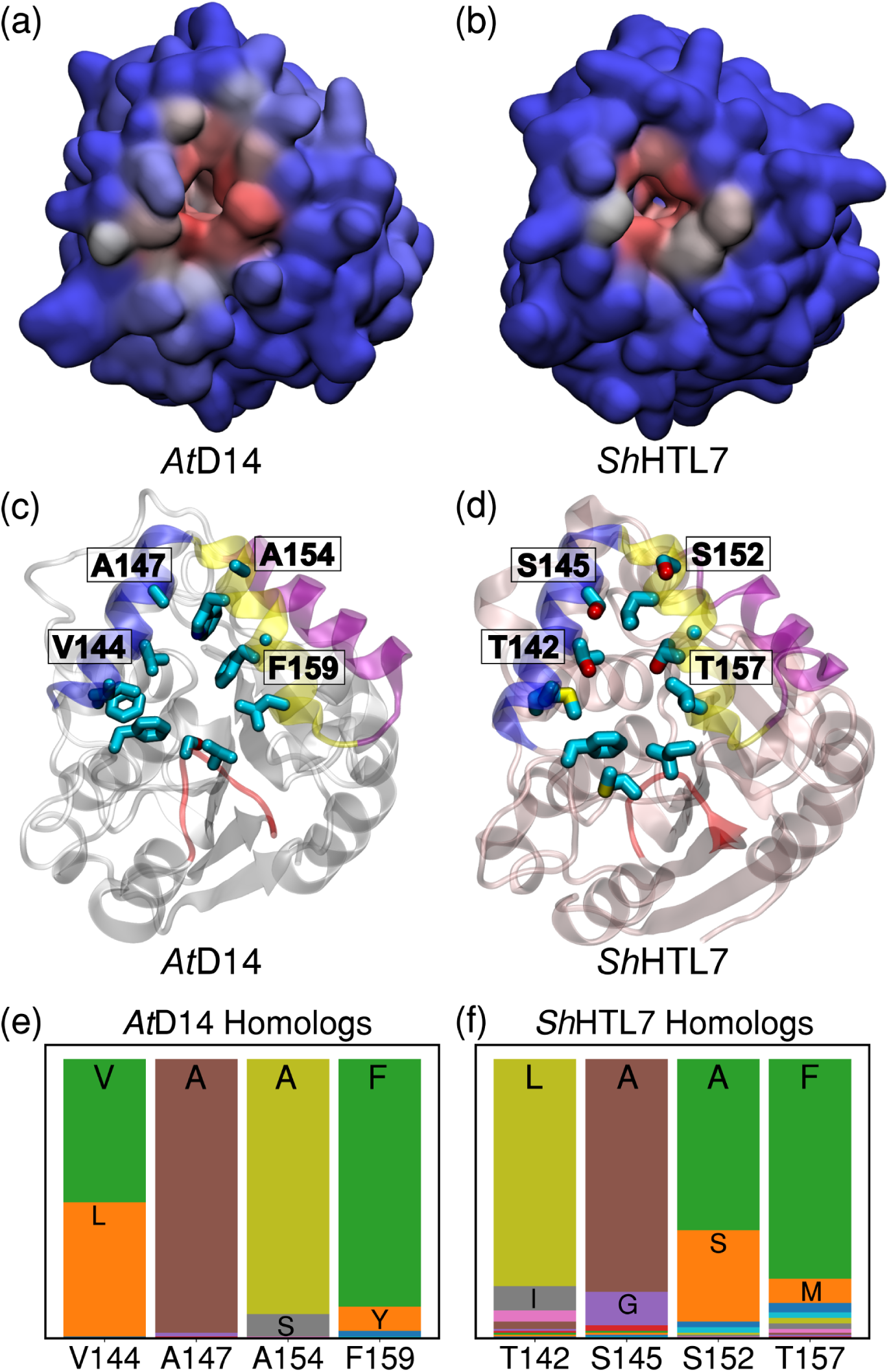
Residue ligand contact probabilities for (a) *At* D14 and (b) *Sh*HTL7. Red regions indicate high ligand contact probability, and blue regions indicate low ligand contact probability. (c) *At* D14 pocket entrance residues in contact with the ligand in the anchored state and (d) corresponding residues in *Sh*HTL7. Hydrophobic to polar substitutions are labeled. (e) Most frequently occurring residues in pocket entrance sites for *At* D14 homologs and (f) *Sh*HTL7 homologs. Residue lists for each site can be found in Table S2.

### Catalytically competent D-loop conformation more stable in *Sh* HTL7 than in *At* D14

Enzymatic hydrolysis of the substrate requires the D-loop of the protein to be in a D-in conformation, in which there is interaction between the aspartate (D218/217) and histidine (H247/246) of the catalytic triad. This is known due to previous mutagenesis experiments that have shown elimination of enzymatic activity upon mutation of any of the catalytic triad residues^7, 12, 39, 40^. We computed free energy profiles of the binding process projected onto catalytic D-catalytic H distance and ligand-pocket distance (Fig. 4). The catalytically active state in which D218/217 is oriented into the binding pocket (D-in), substrate bound state is most stable in *Sh*HTL7, however, *At* D14 exhibits highly stable conformations in which the D218 is oriented away from H247. The

**Fig. 4.**
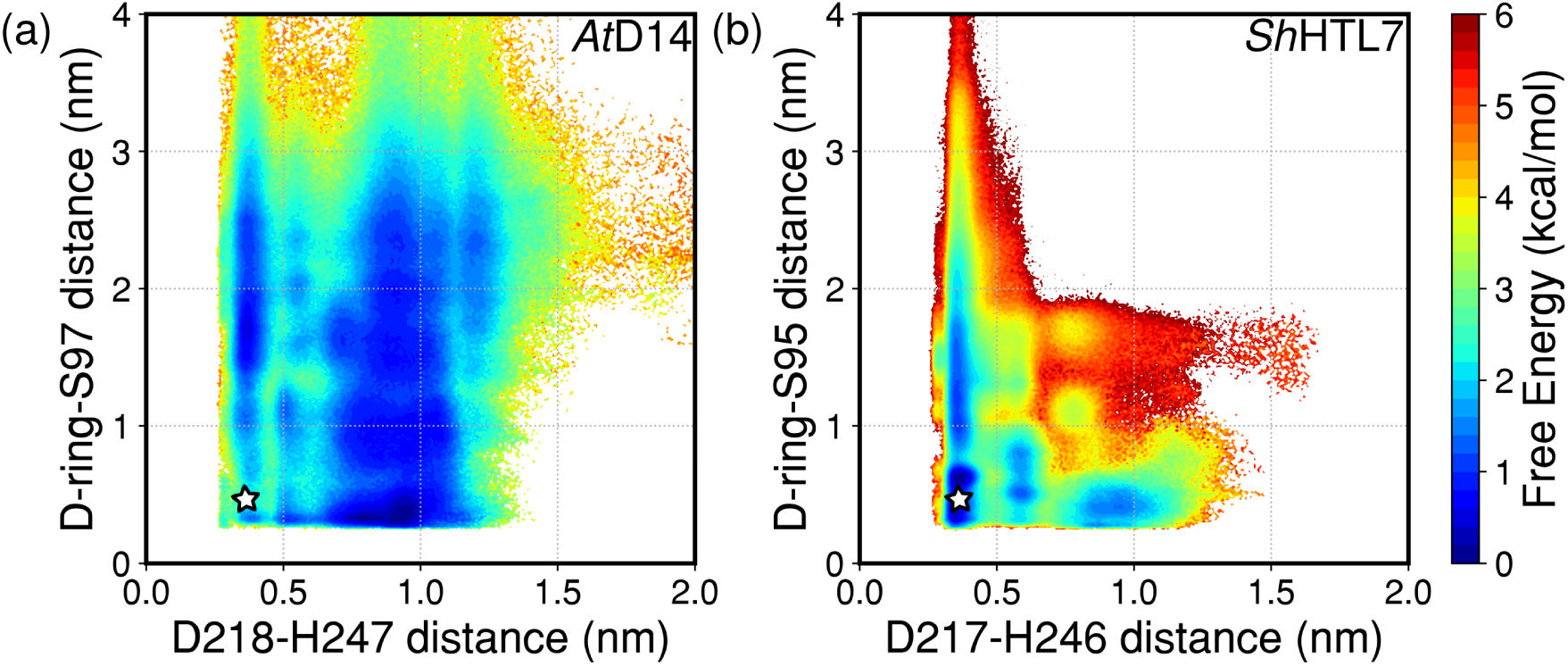
Free energy landscapes of GR24 binding to (a) *At* D14 and (b) *Sh*HTL7 projected onto D218/217-H247/246 distance and ligand-S97/95 distance. The star demarcates the catalytically active state in which the ligand is bound and the aspartate and histidine of the catalytic triad are in contact.

D-out conformations in *At* D14 are *∼*3-5 kcal/mol more stable than in *Sh*HTL7, indicating the presence of stabilizing interactions that facilitate the formation of these catalytically incompetent conformations. Upon comparison of the D-loop sequences in *At* D14 (**AK**DV**S**VPA) and *Sh*HTL7 (**SN**DI**M**VPV), we identified three mutations to differing residue types (i.e. hydrophobic to hydrophilic, charged to neutral): A216S, K217N, and S220M. Based on free energy profiles of key contacts involving these residues, we determine that each of these three mutations contribute to stabilization of the D-out conformation in *At* D14 (Fig. 5).

**Fig. 5.**
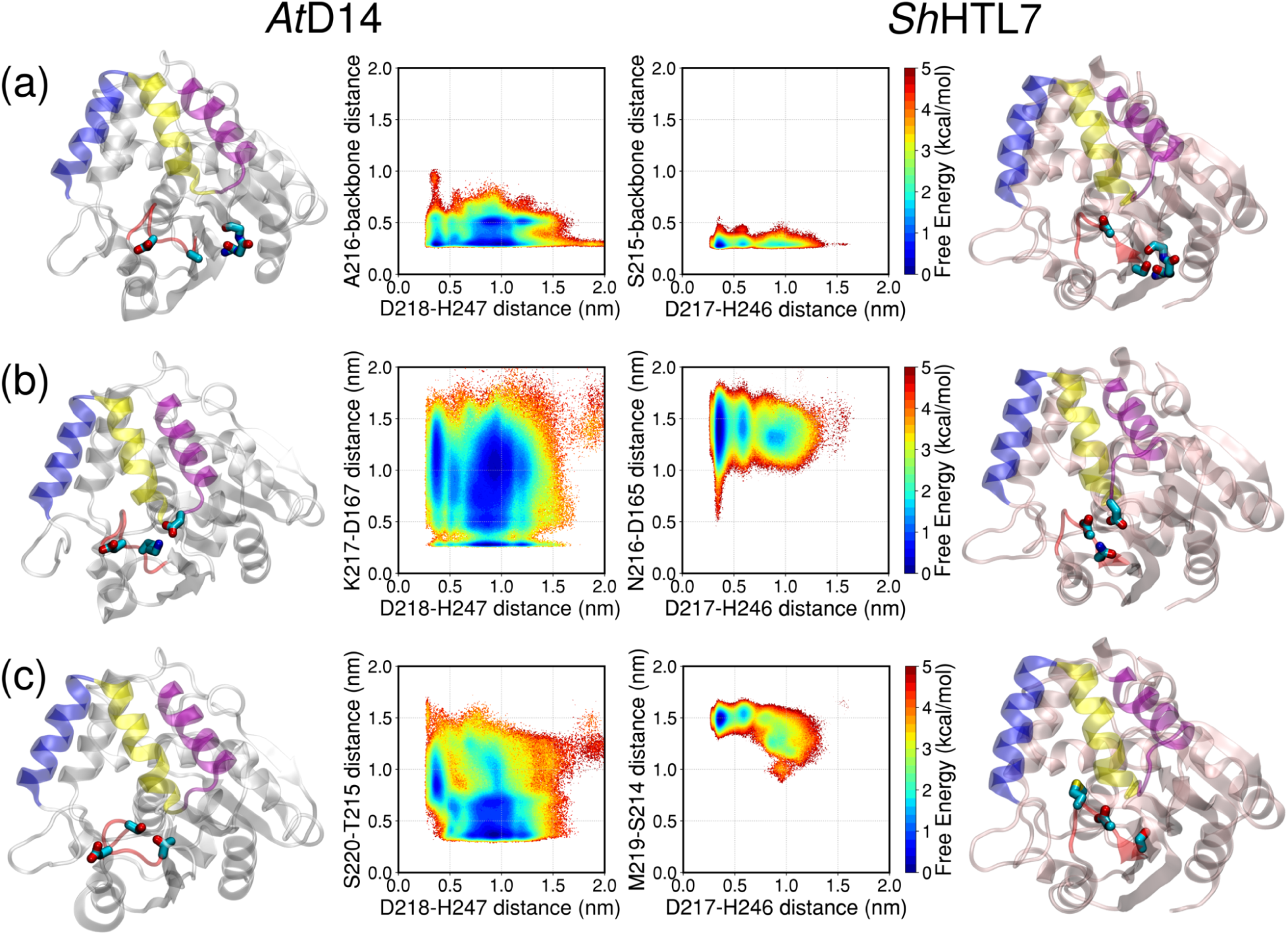
Contacts stabilizing the D-out conformation relative to the D-in conformation in *At* D14. The D-in conformation is defined as having an interaction between D218/217 and H247/246, which means that the D-H distance is within *∼*0.5 nm. (a) The S215-backbone interaction (S215-backbone distance *<*0.5 nm) in *Sh*HTL7 is lost in *At* D14. (b) The K217-D167 salt bridge in *At* D14 (K217-D167 distance *<*0.5 nm) is lost in *Sh*HTL7. (c) The S220-T215 hydrogen bond in *At* D14 (S220-T215 distance *<*0.5 nm) is lost in *Sh*HTL7.

The A216S mutation is located on the end of the D-loop closest to the T2 helix. The corresponding residue in *Sh*HTL7, is S215, which is able to form hydrogen bonding interactions with the adjoining *β*-strand. This limits the range of motion of the D-loop. The A216 residue in *At* D14 is unable to form a hydrogen bond with the adjacent *β*-strand, allowing for increased D-loop motion. Additionally, the free energy landscape indicates that less stable D218-H247 interaction is observed in the absence of A216-backbone interaction (Fig. 5a). This implies that the interaction between S215 in *Sh*HTL7 and the adjacent backbone helps to stabilize the D-loop in the catalytically active D-in conformation.

The K217 residue in *At* D14 can form salt bridges with nearby negatively charged residues (D167, E244). In *Sh*HTL7, the corresponding residue is N216, which eliminates a positive charge and prevents the formation of stable salt bridges. In particular, formation of the K217-E244 salt bridge in *At* D14 destabilizes the D218-H247 interaction, whereas absence of this salt bridge in *Sh*HTL7 allows for a stable D217-H246 interaction (Fig. 5b). In addition to D167, K217 can also form a salt bridge with E244. However, equally stable D218-H247 interactions are observed both in the presence and absence of the K217-E244 salt bridge, which indicates that this interaction does not destabilize the D218-H247 interaction (Fig. S3). In *Sh*HTL7, N216 is able to form a hydrogen bond with E243. As observed in *At* D14, the D217-H246 interaction remains intact both in the presence and absence of the N216-E243 hydrogen bond (Fig. S3).

Finally, S220 in *At* D14 can form a hydrogen bond with T215, which locks D218 in an outward-oriented position. The free energy landscape with respect to D218-H247 distance and S220-T215 distance indicates that D218-H247 contact is nearly eliminated in the presence of the S220-T215 hydrogen bond (Fig 5c). The corresponding residue to S220 in *Sh*HTL7 is M219, which is hydrophobic and thus unable to form a hydrogen bond with S214, the corresponding *Sh*HTL7 residue to T215 in *At* D14. In the absence of a M219-S214 hydrogen bond, a stable D217-H246 interaction is observed.

### Large fluctuations in *At* D14 pocket volume facilitate binding-incapable states and non-productive binding

Previous structural studies have hypothesized that a large binding pocket volume in *Sh*HTL7 is responsible for its uniquely high sensitivity to strigolactones^14, 16, 41^. To evaluate this hypothesis, we computed the probability distributions of pocket volumes in *At* D14 and *Sh*HTL7 over the course of our simulations (Fig. 6). The average pocket volumes of the two proteins are in close agreement with each other (*µ*=268 Å^3^ and 274 Å^3^ for *At* D14 and *Sh*HTL7, respectively). However, *At* D14 displays a significantly broader distribution of binding pocket volumes (*σ*=90 Å^3^ and 45 Å^3^ for *At* D14 and *Sh*HTL7, respectively). Using the same pocket volume calculation metrics, we computed the pocket volumes of the *apo* crystal structures of *At* D14 and *Sh*HTL7 to be 215 Å^3^ and 358 Å^3^, respectively. While the *Sh*HTL7 crystal structure does have a larger binding pocket volume than the *At* D14 crystal structure, this difference in pocket volume decreases significantly in an aqueouse environment. In both proteins, the primary modulator of binding pocket volume is a hinging motion between the T1 and T2 helices (Fig. S4). In the lowest-volume states, the binding pocket becomes solvent-inaccessible, rendering the protein unable to bind ligand. The highest-volume states allow for a large ensemble of ligand binding poses to form including many non-productive binding states in which the ligand is inside the pocket but not positioned for hydrolysis. These results indicate that the decreased tendency of *Sh*HTL7 to change its pocket volume may serve to increase its catalytic efficiency by retaining the pocket in a solvent-accessible conformation while also decreasing the stability of non-productive binding poses.

**Fig. 6.**
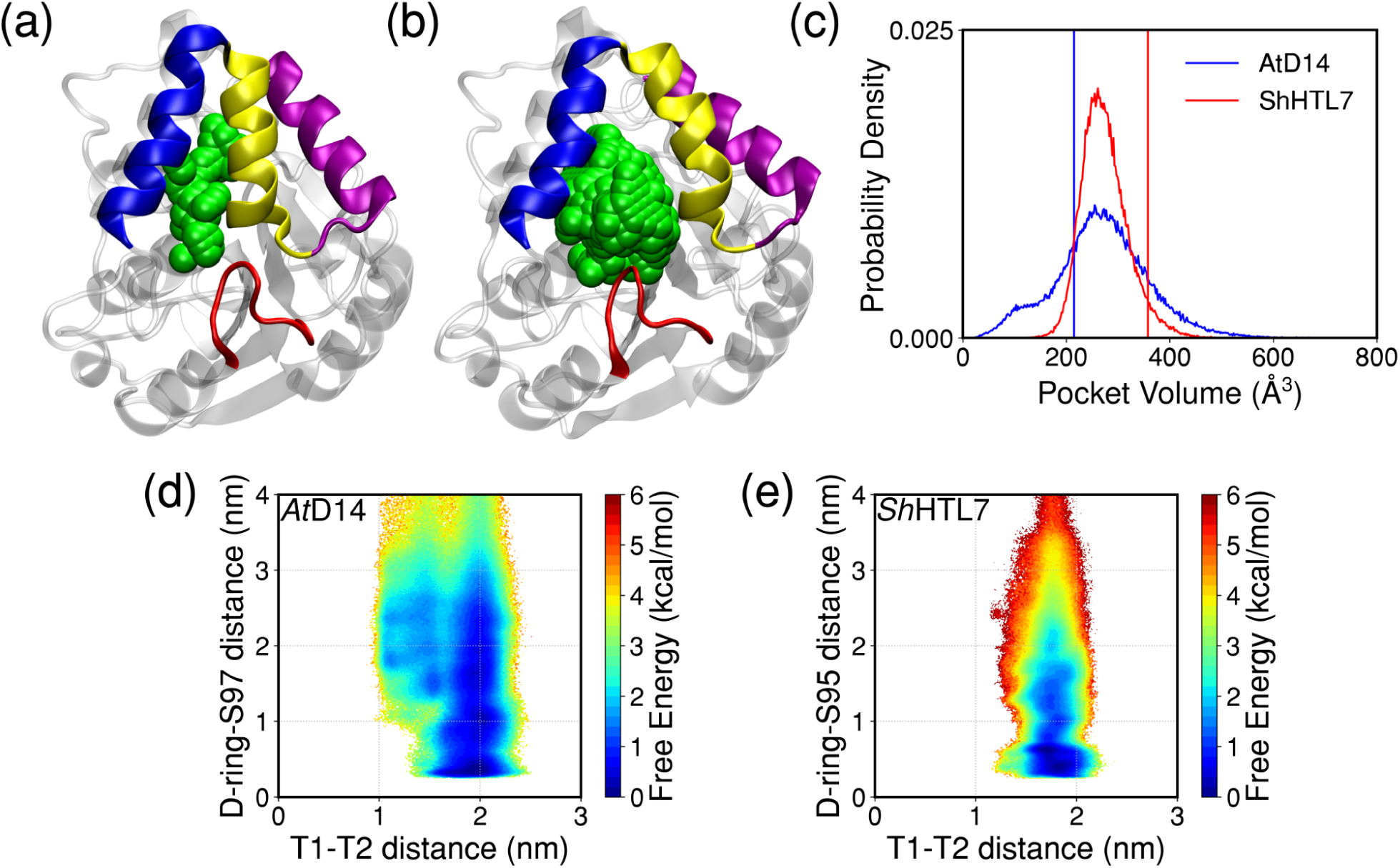
(a) Representative low-volume structure of *At* D14 observed in simulations (b) Representative high-volume structure of *At* D14 observed in simulations. (c) Probability distributions of pocket volume for *At* D14 and *Sh*HTL7. Vertical lines indicate the crystal structure pocket volumes. (d) Free energy landscape of ligand binding and T1-T2 distance for *At* D14. (e) Free energy landscape of ligand binding and T1-T2 distance for *Sh*HTL7.

The low-volume states accessible by *At* D14 are stabilized by hydrophobic interactions between the T1 and T2 helices, indicating that polarity in this region may play a dual role in modulating ligand selectivity. As previously stated, these residues enable a stable intermediate state to form, which acts as a barrier to binding. The low-volume states in which the pocket is nearly solvent inaccessible also show hydrophobic interactions between T1 and T2 helix residues, which indicates that these interactions could play a role in stabilizing low-volume states as well. In addition to the hydrophobic contacts between the T1 and T2 helices, a non-conserved salt bridge between the T1 and T4 helices in *At* D14 provides stabilization to the low-volume states as well (Fig. 7). Both *At* D14 and *Sh*HTL7 have a conserved arginine on the T4 helix (R192/191) that are able to form a salt bridge with E142/E140 on the T1 helix. However, *At* D14 also has a second negative residue, E138, on T1 that can form a salt bridge with R192. The free energy landscape of this interaction and the T1-T2 distance indicates that in the presence of this E138-R192 salt bridge, low volume states are stabilized. This residue is mutated to Q136 in *Sh*HTL7, so a salt bridge cannot form and stabilize low-volume states of the protein.

**Fig. 7.**
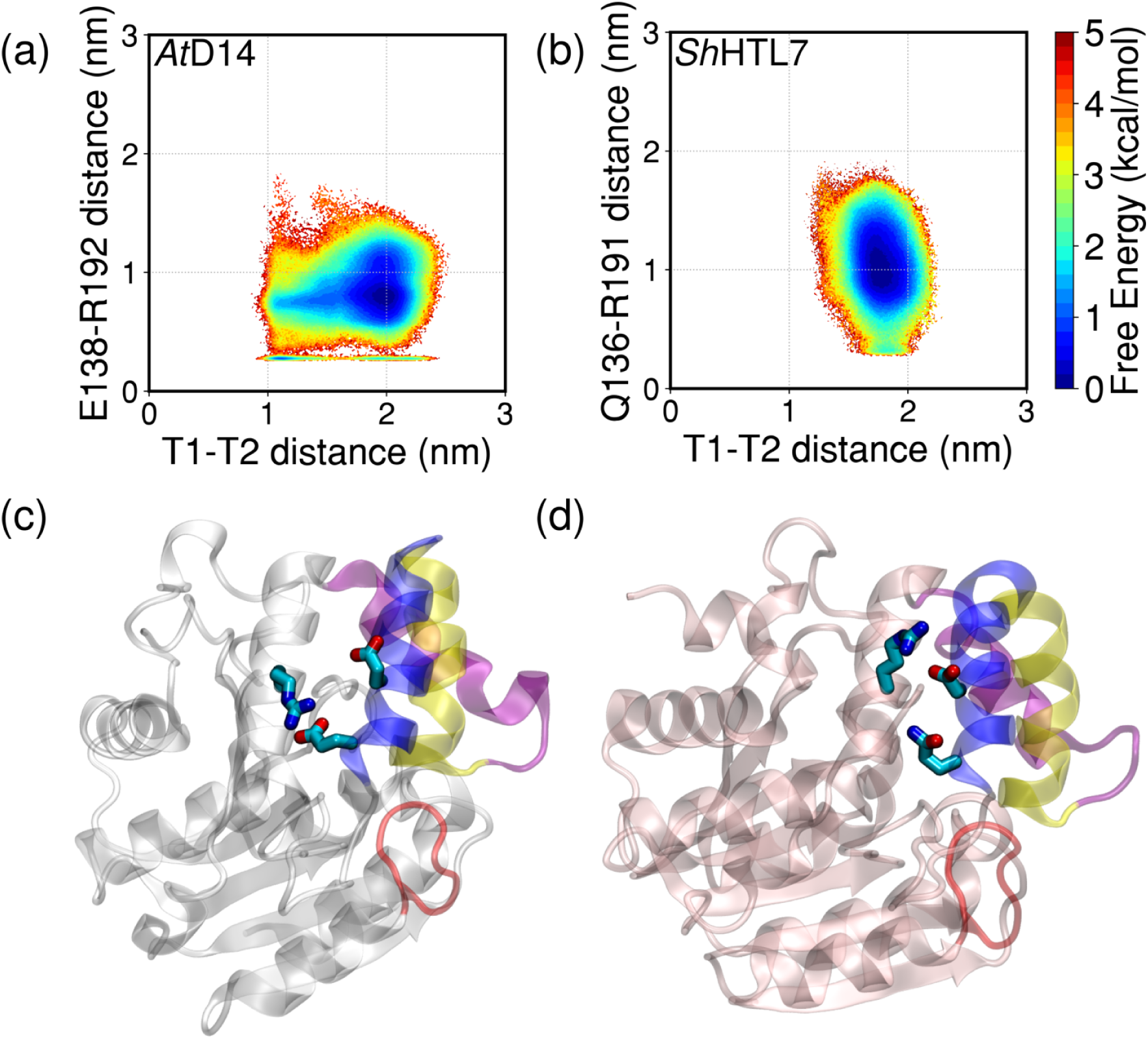
Additional salt bridge in *At* D14 which stabilizes the low-volume states. (a) Free energy landscape showing stabilization of low-volume states in the presence of the E138-R192 salt bridge in *At* D14, (b) *Sh*HTL7 free energy landscape showing little Q136-R191 interaction as well as lower population of low-volume states than in *At* D14. (c) E138-R192 salt bridge in *At* D14. E142, which can also form a salt bridge with R192, is also shown. (d) *Sh*HTL7 salt bridge between R191 and D140, the corresponding residue to E138 in *At* D14. Q136, the corresponding residue to E138 in *At* D14, is also shown.

## 3 Discussion

Using extensive MD simulations, we have characterized the mechanism of substrate binding to strigolactone receptors in full atomistic detail and identified several key differences in the binding mechanisms that contribute to the ligand selectivity between strigolactone receptors. Based on our simulations, GR24 binds to both *At* D14 and *Sh*HTL7 in the same binding pose as the reported crystal structure of *Os*D14-GR24 complex^18^. Additionally, since our simulations were performed in an unbiased manner, we were also able to identify several non-productive bound states in which the ligand is bound but improperly positioned for hydrolysis.

In addition to characterizing the possible ligand poses within the receptor binding pockets, we identified a key anchored intermediate along the binding pathway that is *∼*1.5 kcal/mol more stable in *At* D14 than in *Sh*HTL7. This difference in stability indicates that the anchored state population relative to the bound state is *∼*12 times higher in *At* D14 than in *Sh*HTL7. This likely results in faster binding kinetics in *Sh*HTL7 than in *At* D14, which in turn would lead to a higher observed catalytic turnover rate. Since hydrolysis is driver of receptor activation and downstream signaling, faster binding kinetics can contribute to enhanced signaling in *Sh*HTL7 compared to *At* D14.

We also identified several key interactions involving residues on the D-loop of *At* D14 that stabilize the D-loop in a catalytically inactive, D218-out conformation. While these interactions do not preclude binding, they likely hinder the catalytic process in *At* D14 compared to in *Sh*HTL7 by stabilizing catalytically inactive states of the protein. Assuming that the hydrolysis is an inducer of receptor activation and downstream signaling, the stabilization of catalytically inactive conformations of the protein can lead to decreased signaling as well.

Finally, we evaluated the hypothesis that a larger binding pocket in *Sh*HTL7 enables its unique sensitivity to strigolactone. The average binding pocket volumes of *At* D14 and *Sh*HTL7 are nearly identical, however, *At* D14 is able to access more low-volume states which preclude ligand binding as well as more high-volume states which allow more non-productive binding to occur. These effects play a dual role in modulating ligand selectivity: *Sh*HTL7 is less likely to adopt low-volume states which are unable to bind ligand as well as high-volume states that are likely to bind ligand in non-productive, signaling-inactive poses.

A 10000-fold difference in EC50 for downstream signaling implies a difference of *∼*5.5-6 kcal/mol along the strigolactone signaling process. In total, combined effects of enhanced binding and more stable catalytically active state contribute *∼*2-3 kcal/mol. This indicates that while the differences in the ligand binding process are contributors to the unique sensitivity of *Sh*HTL7, subsequent steps in the signaling process, such as hydrolysis, activation, and association with signaling partners, play important roles in modulating this selectivity as well.

To our knowledge, this is the first in-depth characterization of the ligand binding process in strigolactone receptors. This study demonstrates the utility of molecular simulations approaches in providing mechanistic insights into fundamental questions in the field of plant biology^42, 43^. Due to the importance of strigolactone signaling in crop productivity and parasitic weed germination, there is great interest in developing strigolactone signalling antagonists^44–47^. The factors we have identified that modulate ligand selectivity in strigolactone receptors can be used to inform the design of selective signalling agonists to enhance shoot branching in crops, induce suicidal germination in parasitic weeds, or prevent parasitic weed germination.

## 4 Methods

### Molecular dynamics simulations

MD simulations were prepared using AmberTools 14/18 and run using Amber 14/18^48^. The protein was described using the ff14SB force field and water was described using the TIP3P model^49^. The GR24 ligand was described using the generalized AMBER force field (GAFF)^50^. Force field parameters for GR24 were generated using Antechamber. Initial structures for *At* D14 and *Sh*HTL7 were obtained from Protein Data Bank entries 4IH4^51^ and 5Z7Y^16^, respectively. The GR24 substrate was superimposed into the binding pocket by structural alignment of the bound structure of *Os*D14 bound to GR24 (PDB 5DJ5)^18^. For *Sh*HTL7, an additional system was prepared with the ligand randomly placed in solution using Packmol^52^. Each protein-ligand complex was solvated in a TIP3P water box of size *∼*70 *×* 70 *×* 70 Å. NaCl was added at a concentration of 0.15M to neutralize the system. Each structure was minimized for 10000 steps using the conjugate gradient descent method and equilibrated for 10 ns. Production runs were performed for an aggregate of 207 *µ*s for *Sh*HTL7 and 198 *µ*s for *At* D14. Temperatures were held constant at 300 K using the Berendsen thermostat, and pressures were held constant at 1.0 bar using the Berendsen barostat. Full electrostatics were computed using the Particle Mesh Ewald algorithm with a cutoff distance of 10 Å^53^. Bonds to hydrogen were constrained using the SHAKE algorithm^54^.

### Markov state model construction

Markov state models (MSMs) were constructed using the PyEmma^55^ package. Thirty-one input distance features were computed using MDTraj 1.9.0^56^ (Table S3). The input distances were projected onto a reduced set of coordinates using time-lagged independent component analysis (TICA)^57, 58^. The dimensionality-reduced coordinates were then clustered into states using the Mini-Batch K-Means algorithm prior to MSM estimation. The hyperparameters (number of TICA dimensions and number of clusters) were chosen via maximization of cross-validation scores (Fig. S6)^59^. Lag time was chosen by convergence implied timescales with respect to lag time (Fig. S5). Final parameters for MSM construction are listed in Table S4. Markovianity of the model was further validated using the Chapman-Kolmogorov test (Fig. S8). Free energy landscapes were calculated by computing the probability distribution along chosen sets of order parameters (*x, y*) and weighting each point by the equilibrium probability of its associated MSM state (Eq. 1).

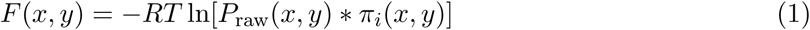

### Standard binding free energy calculation

Standard free energies of binding were calculated using the volume correction method as detailed in Buch *et al*.^27^. Briefly, this method corrects for non-standard ligand concentration in the simulation by introducing a correction term (Eq. 2) that corresponds to the free energy of moving the ligand from a 1M solution to simulation conditions. For calculation of bound state volume, the bound state was defined as points within 1.0 nm and 4.0 kcal/mol of the minimum free energy point on the 3-dimensional MSM-weighted free energy landscape projected onto ligand position. Convergence of Δ*G*_0_ with respect to bound state definition is shown in Fig. S9.

### Binding pocket volume calculation

Binding pocket volumes were calculated using the POVME 2.0 package^60^. For each protein, a “maximum englobing region” was defined as a sphere centered at the midpoint between the geometric center of the T1 and T2 helix C-*α* atoms (residues 138-165 for *At* D14, residues 136 to 163 for *Sh*HTL7) and the C-*α* atom of the catalytic serine (Fig. S10). The radius of the maximum englobbing sphere was set as the distance between the center and the C-*α* of the catalytic serine. The probability distribution of binding pocket volumes was weighted by Markov state model equilibrium probability. The average and standard deviation of the pocket volumes were calculated using the MSM-weighted probability distributions.

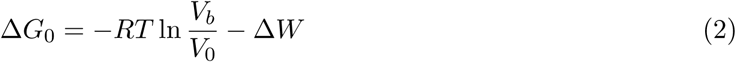

### Residue-ligand contact probability calculation

Residue-ligand distances for each residue were computed using MDTraj 1.9.0^56^. Contacts were defined as residue-ligand distances within a cutoff distance of 4.0 Å. The equilibrium contact probability was calculated as the product of raw contact probability within each MSM state multiplied by the equilibrium probability of the MSM state as shown in Eq. 3:

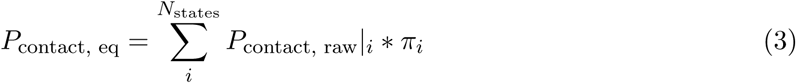

## 5 Acknowledgements

This research was supported by the Blue Waters sustained-petascale computing project, which is funded by the National Science Foundation (OCI-0725070 and ACI-1238993) and the state of Illinois. J.C. acknowledges support from the Samuel W. Parr Graduate Fellowship (Department of Chemical and Biomolecular Engineering, University of Illinois) and the National Institutes of Health Chemistry-Biology Interface Training Grant (T32-GM070421). D.S. acknowledges support from the Center for Advance Study at University of Illinois at Urbana-Champaign.

## Notes

### Competing Interest Statement

The authors have declared no competing interest.

