## Supplementary Information for "Role of substrate recognition in modulating strigolactone receptor selectivity in witchweed"

### Supporting Information for: Role of substrate recognition in modulating strigolactone receptor selectivity in witchweed

Jiming Chen[1], Alexandra White[2], David C. Nelson[2], and Diwakar Shukla[1,3,4,5,6*]

[1]*Department of Chemical and Biomolecular Engineering, University of Illinois at Urbana-Champaign, Urbana, IL 61801*
[2]*Department of Botany and Plant Sciences, University of California, Riverside, Riverside, CA 92521*
[3]*Center for Biophysics and Quantitative Biology, University of Illinois at Urbana-Champaign, Urbana, IL 61801*
[4]*National Center for Supercomputing Applications, University of Illinois at Urbana-Champaign, Urbana, IL 61801*
[5]*Beckman Institute for Advanced Science and Technology, University of Illinois at Urbana-Champaign, Urbana, IL 61801*
[6]*NIH Center for Macromolecular Modeling and Bioinformatics, University of Illinois at Urbana-Champaign, Urbana, IL 61801*
***

#### 1 Sequence and secondary structure alignments of *At*D14 and *Sh*HTL7

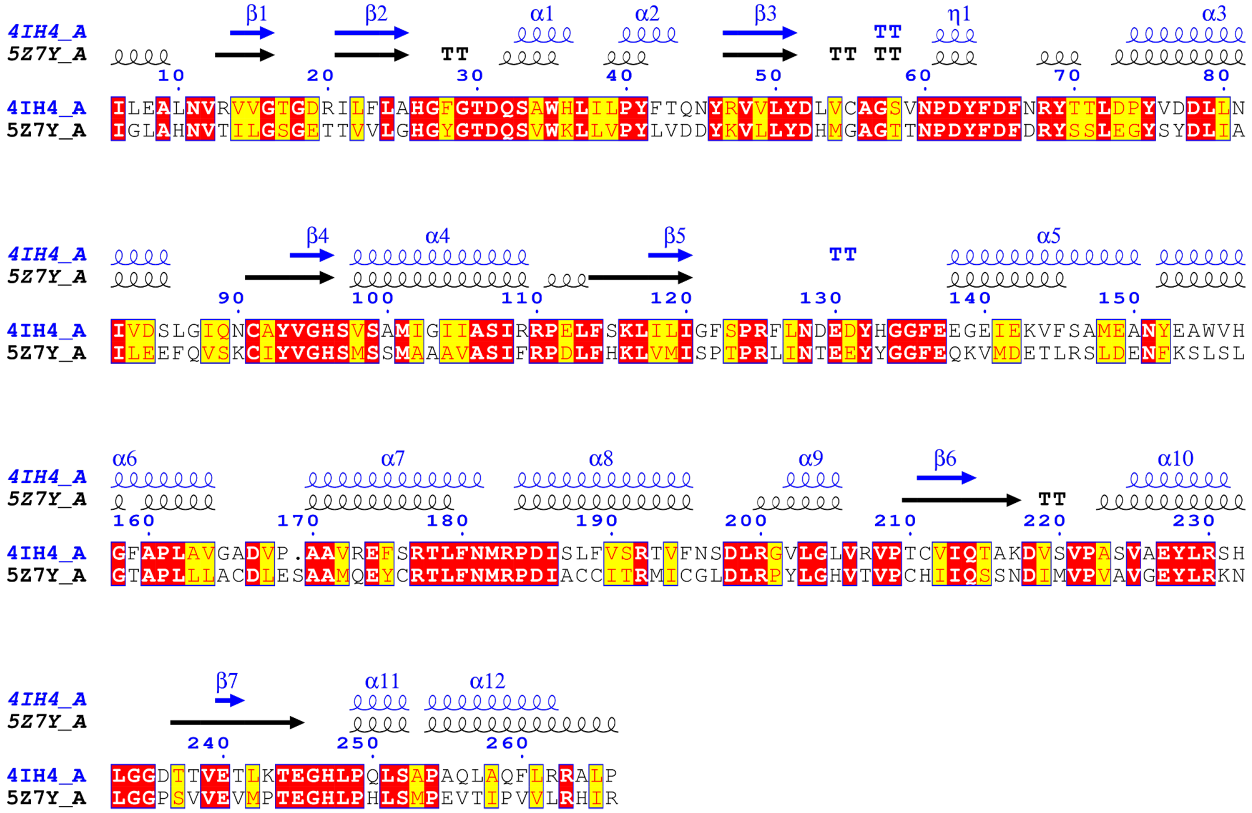


**Figure S1.** Sequence and secondary alignment of *At*D14 and *Sh*HTL7 based on chain A of 4IH4 (*At*D14) and 5Z7Y (*Sh*HTL7). The sequences share ∼44% identity and high conservation of secondary structure. This alignment was generated using the EScript 3.0 web server. The $\alpha$5-$\alpha$8 helices in the alignment correspond to the T1-T4 helices.

#### 2 Free energy landscape over slowest processes

To further investigate the free energy profiles of the binding processes, we projected the free energy onto the two slowest time-lagged independent components (TICs) of each system (Fig. S2). TICs are linear combinations of the input features (Table S3) that produce the maximum time-lagged autocorrelation of a set of time-series data. This identifies a set of order parameters for a system that correspond to the longest time-scale processes captured by the simulations. For comparison, we projected the data for both systems onto the TICs computed with both *At*D14 simulations and *Sh*HTL7 simulations.

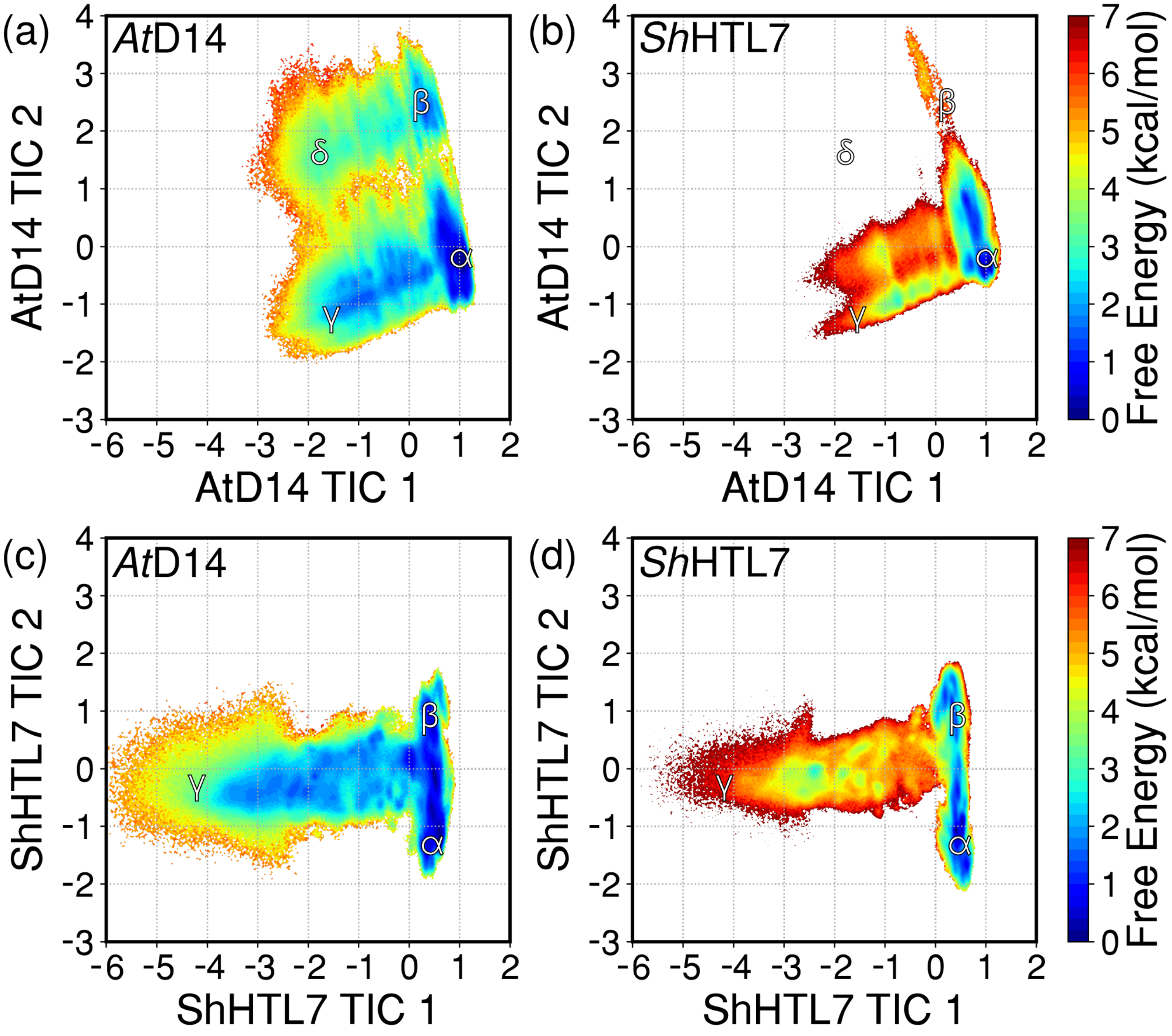


**Figure S2.** Free energy landscapes of (a) *At*D14 binding projected onto *At*D14 TICs, (b) *Sh*HTL7 binding projected onto *At*D14 TICs, (c) *At*D14 binding projected onto *Sh*HTL7 TICs, and (d) *Sh*HTL7 binding projected onto *Sh*HTL7 TICs. Labeled states for *At*D14 TICs (a,b) are $\alpha$: Bound, D-in, $\beta$: Bound, D-out, $\gamma$: Unbound, D-in, and $\delta$: Unbound, D-out. Labeled states for *Sh*HTL7 TICs (c,d) are $\alpha$: Bound, $\beta$: Inverse bound, $\gamma$: Unbound.

#### 3 Calculation of site conservation

Site conservation scores of key residues at the pocket entrance were calculated using the ConSurf webserver. Scores are normalized such that the average score for all residues is zero and the standard deviation is one, and more negative scores indicate higher conservation. Settings used for calculation of scores are listed in Table S1.

| | *At*D14 | *Sh*HTL7 |
|---|---|---|
| PDB ID | 4IH4 | 5Z7Y |
| Multiple Sequence Alignment | MAFFT | MAFFT |
| Homolog Source | UNIREF90 | UNIREF90 |
| Homolog Search Algorithm | HMMER | HMMER |
| HMMER E-value | 0.0001 | 0.0001 |
| HMMER Iterations | 1 | 1 |
| Maximal % Identity | 95 | 95 |
| Minimal % Identity | 50 | 50 |
| Number of Sequences | 150 | 150 |

**Table S1.** Parameters used for calculation of site conservation

Conservation scores and most frequent residues for pocket entrances sites are listed in Table S2.

| Site | Conservation Score | Most Frequent Residues |
|---|---|---|
| *At*D14 Homologs | | |
| V144 | -0.422 | V,L,F |
| A147 | -0.819 | A,G,R |
| A151 | -0.505 | A,S,E |
| F159 | -0.030 | F,Y,W,H |
| *Sh*HTL7 Homologs | | |
| T142 | 0.115 | L,I,V,M,X,W,T,F,A |
| S145 | -0.305 | A,G,S,X,M,L,E |
| S149 | -0.074 | A,S,V,T,L,E,D |
| T157 | 0.477 | F,M,L,Y,W,S,T,V,R,H |

**Table S2.** Conservation scores and most frequent residues for key pocket entrance residues. The most frequently appearing residues are listed in decreasing order of frequency.

#### 4 Additional K217 salt bridges in *At*D14

Free energy landscapes show that D218/217-H247/246 interactions remain intact in the presence of K217-E244, N216-E243 interactions (Fig. S3) This is another salt bridging interaction that is present in *At*D14 but not *Sh*HTL7, but unlike the other D-loop interactions, this particular salt bridge has little effect on D218-H247 interaction.

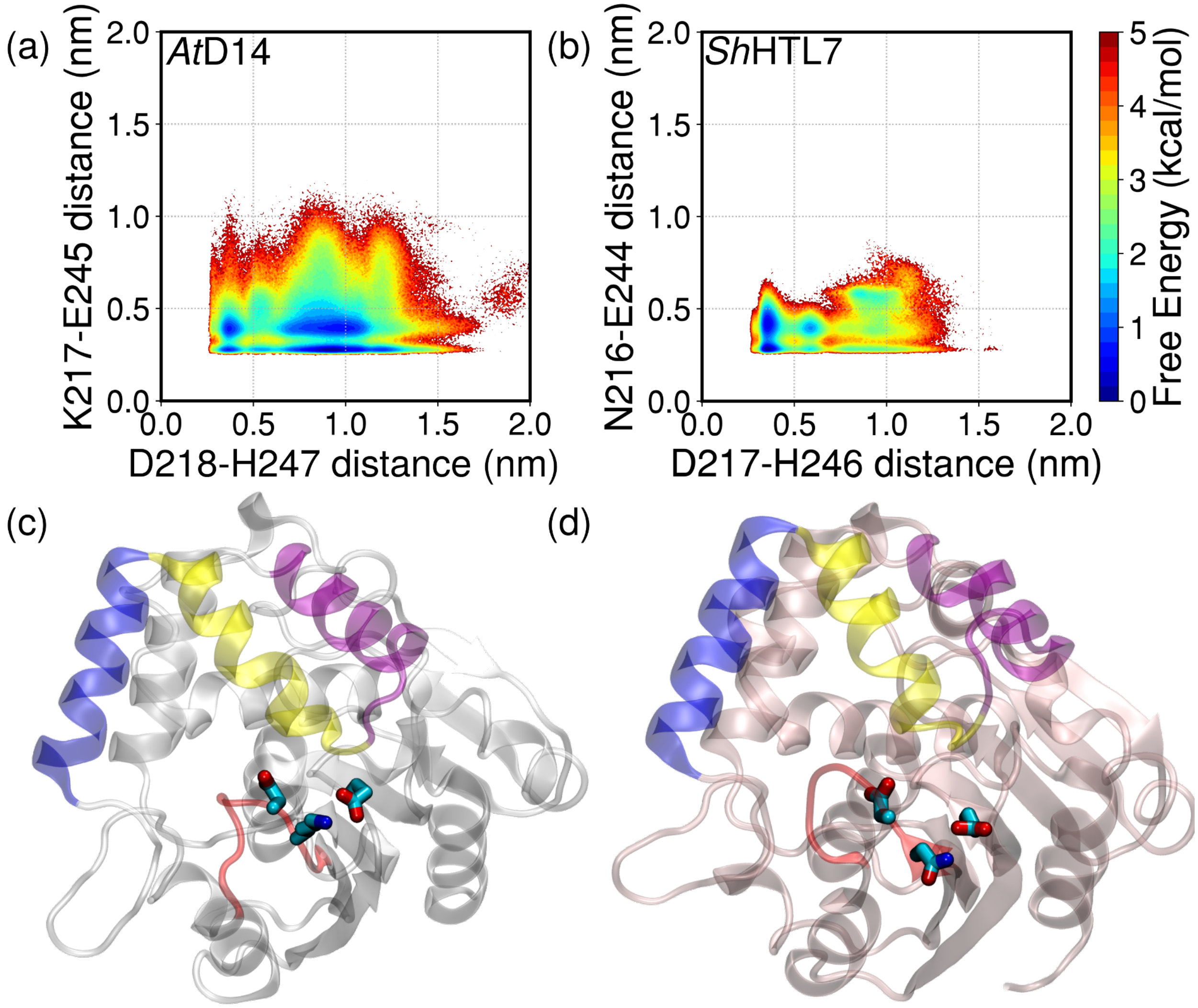


**Figure S3.** A K217-E244 salt bridge can form in *At*D14 that does not destabilize the D218-H247 interaction

#### 5 T1-T2 motion modulates binding pocket volume

Plots of pocket volume vs. T1-T2 distance are shown in Fig. S4. Low-volume states are accessed by a hinging motion of the T1 and T2 helices. T1-T2 distances were calculated using the C-$\alpha$ atoms of E140 and L162 for *At*D14 and V138 and L160 for *Sh*HTL7.

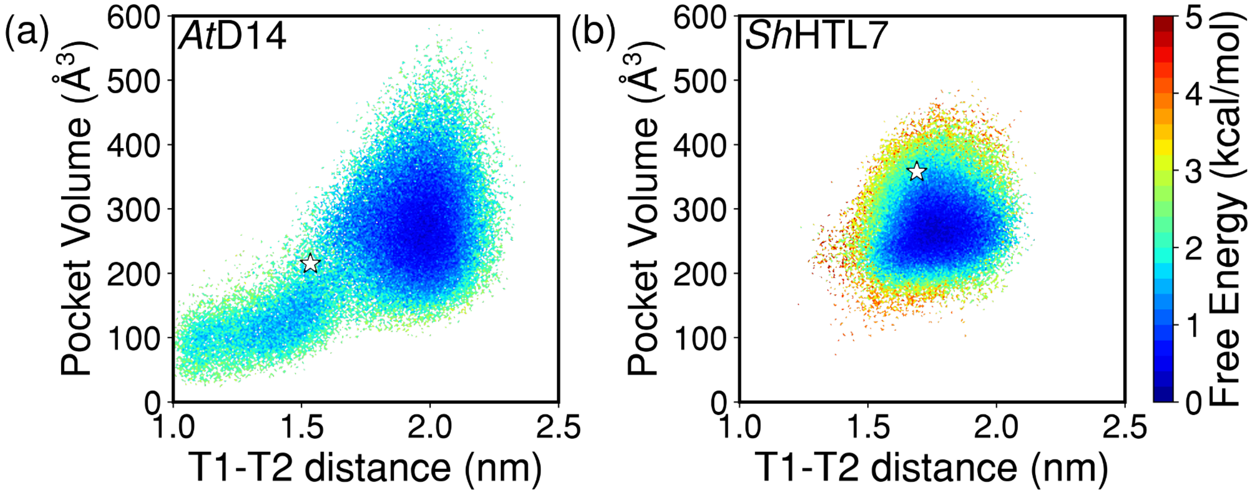


**Figure S4.** Pocket volume vs. T1-T2 distance for (a) *At*D14 and (b) *Sh*HTL7. Stars denote the pocket volume and T1-T2 distances of the crystal structures.

#### 6 Markov state model construction and validation

**Distance features used for MSM construction.** Table S3 lists the 30 distance features used for MSM construction. All protein-ligand distances were calculated using closest heavy-atom distance, and all protein-protein distances were calculated using C-$\alpha$ atom distances.

| | *At*D14 | *Sh*HTL7 |
|---|---|---|
| Ligand-catalytic S distances | A-ring-S97 | A-ring-S95 |
| | D-ring-S97 | D-ring-S95 |
| T1-T2 distances | M148-W155 | L146-L153 |
| | V144-F159 | T142-T157 |
| | E140-L162 | V138-L160 |
| A-ring-T1 distances | A-ring-A141 | A-ring-M139 |
| | A-ring-V144 | A-ring-T142 |
| | A-ring-A147 | A-ring-S145 |
| A-ring-T2 distances | A-ring-W155 | A-ring-L153 |
| | A-ring-F159 | A-ring-T157 |
| | A-ring-L162 | A-ring-L160 |
| D-ring-T1 distances | D-ring-A141 | D-ring-M139 |
| | D-ring-V144 | D-ring-T142 |
| | D-ring-A147 | D-ring-S145 |
| D-ring-T2 distances | D-ring-W155 | D-ring-L153 |
| | D-ring-F159 | D-ring-T157 |
| | D-ring-L162 | D-ring-L160 |
| D-loop-catalytic H distances | A216-H247 | S216-H246 |
| | K217-H247 | N216-H246 |
| | D218-H247 | D217-H246 |
| | V219-H247 | I218-H246 |
| | S220-H247 | M219-H246 |
| | V221-H247 | V220-H246 |
| | P222-H247 | P221-H246 |
| A-ring-D-loop distances | A-ring-D218 | A-ring-D217 |
| | A-ring-S220 | A-ring-M219 |
| | A-ring-P222 | A-ring-P221 |
| D-ring-D-loop distances | D-ring-D218 | D-ring-D217 |
| | D-ring-S220 | D-ring-M219 |
| | D-ring-P222 | D-ring-P221 |

**Table S3.** Featurizations used for MSM construction

**Convergence of implied time scales.** Lag times for the MSMs were chosen by convergence of implied timescales with respect to lag time. In a Markovian system, the implied timescales of the slowest processes should be independent of the lag time used to construct a MSM. Fig. S5 shows convergence of the ten slowest timescales with increasing lag times. A lag time of 20 ns was chosen for both *At*D14 and *Sh*HTL7.

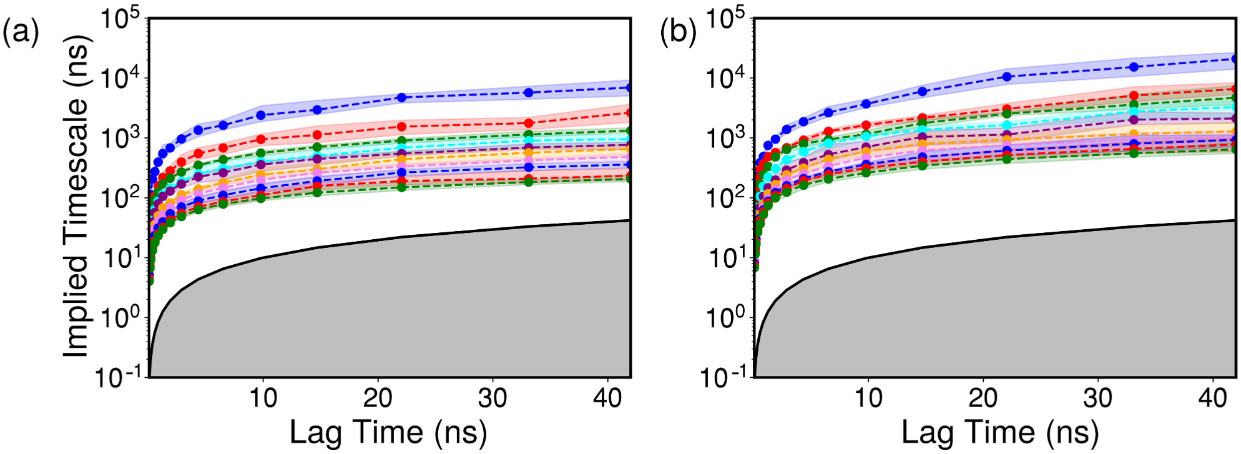


**Figure S5.** Implied time scale plots for (a) *At*D14 and (b) *Sh*HTL7.

**Optimization of MSM hyperparameters.** Hyperparameters for each MSM were chosen by shuffle-split cross-validation scores of models built with TICA components ranging from 2 to 10 and clusters ranging from 100 to 500 (Fig. S6). Models for each set of parameters were cross-validated for 10 trials. Final parameters for MSM construction are listed in Table S4.

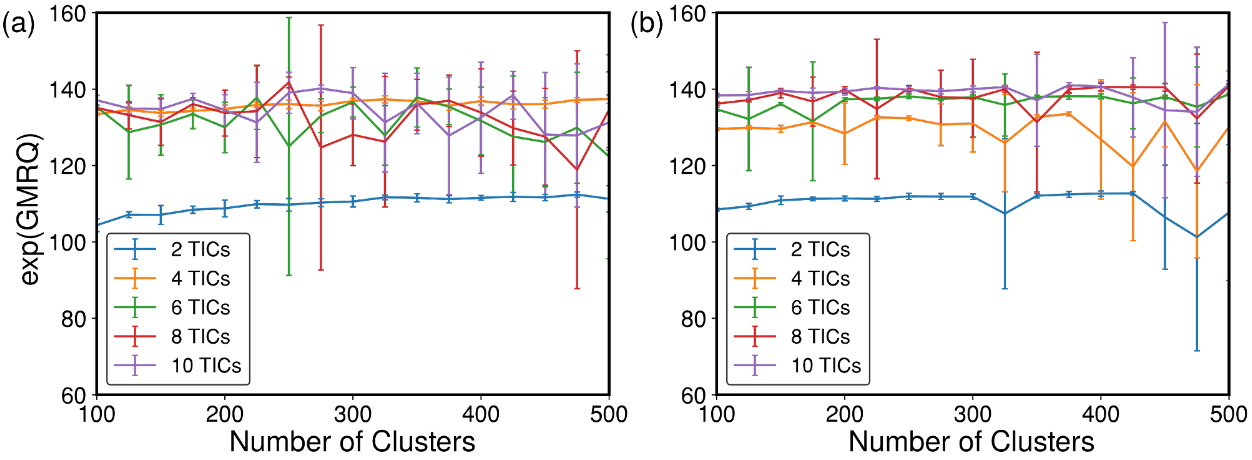


**Figure S6.** Cross-validation scores for (a) *At*D14 and (b) *Sh*HTL7.

| | *At*D14 | *Sh*HTL7 |
|---|---|---|
| Lag time (ns) | 20 | 20 |
| Number of TICA components | 4 | 10 |
| Number of clusters | 325 | 375 |

**Table S4.** Parameters used for MSM construction

**Chapman-Kolmogorov validation.** Further validation of the Markov state models was performed via the Chapman-Kolmogorov test. Briefly, this test shows Markovian behavior of the system by checking that MSM transition matrices estimated at lag time $n\tau$ are equal to the transition matrix estimated at lag time $\tau$ multiplied by itself $n$ times (Eq. S1). A Chapman-Kolmogorov computed on the full (325 states for *At*D14, 375 states for *Sh*HTL7) transition matrices (Fig. S7) shows good agreement between estimated $(T_{ij}(n\tau))$ and predicted $(T_{ij}(\tau)^n)$ transition matrices.

$$T_{ij}(n\tau) = T_{ij}(\tau)^n \qquad (1)$$

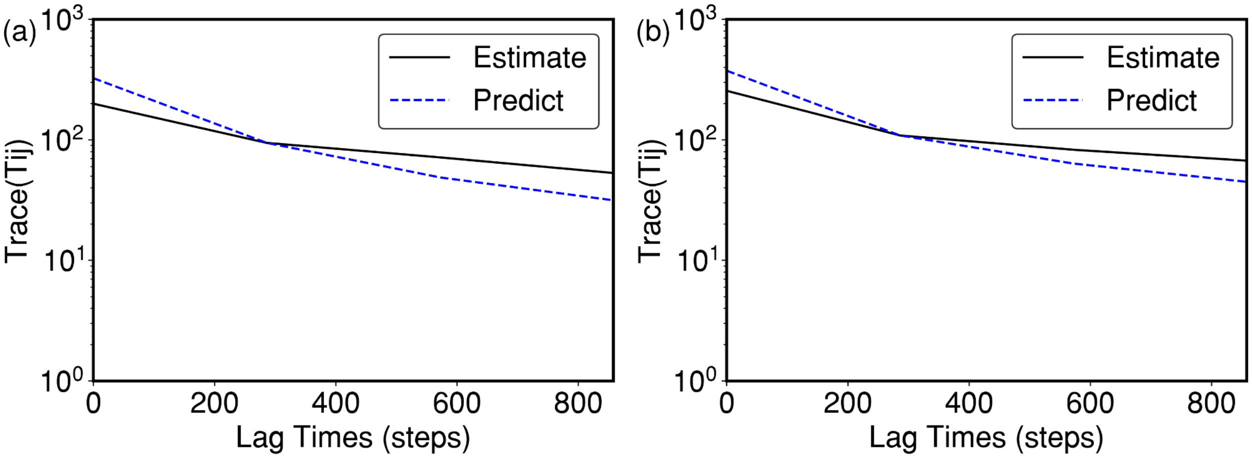


**Figure S7.** Chapman-Kolmogorov tests on $T_{ij}$ showing good agreement between the trace of $T_{ij}$ for (a) *At*D14 and (b) *Sh*HTL7.

In addition to the Chapman-Kolmogorov test on the full transition matrix $T_{ij}$, we also performed a Chapman-Kolmogorov test on five metastable states (Fig. 8). This test shows excellent agreement between estimated and predicted coarse-grained transition matrices.

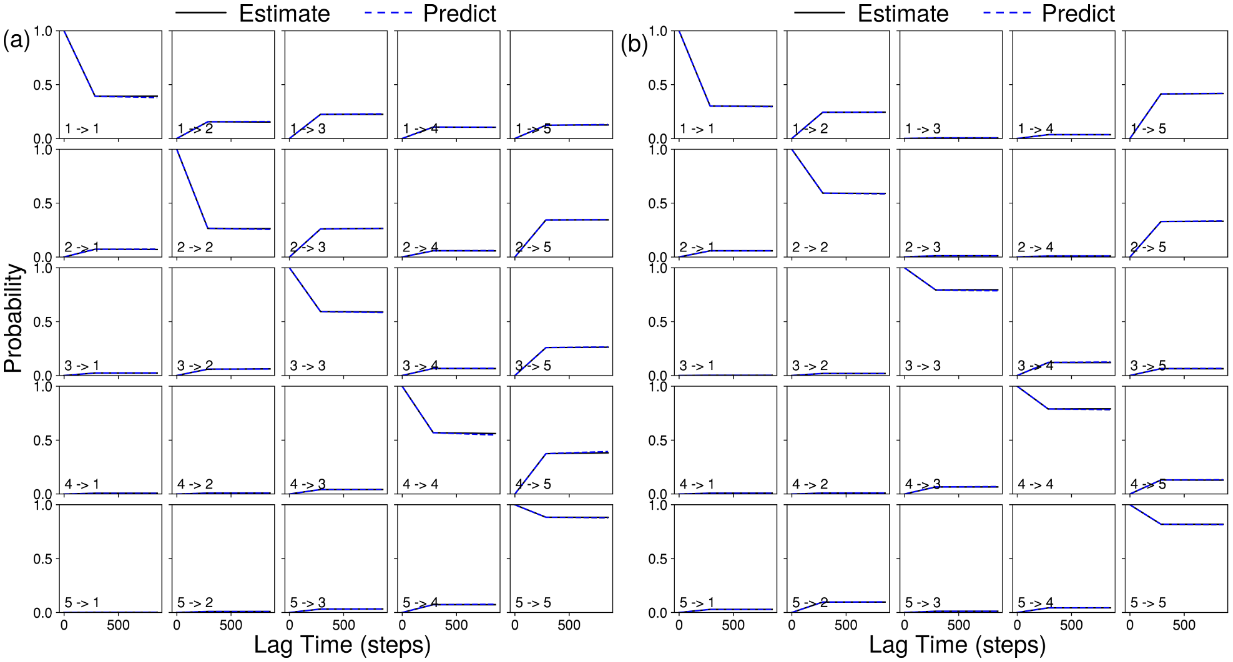


**Figure S8.** Chapman-Kolmogorov tests on five metastable states showing Markovian behavior for (a) *At*D14 and (b) *Sh*HTL7.

#### 7 Cutoff parameters for binding free energy calculation

To calculate the standard binding free energy using the volume correction method, we defined the bound volume on the 3-dimensional free energy landscape as points within 1 nm and 4 kcal/mol of the bound minimum. The spatial constraint was introduced to prevent unbound minima from being included in the bound volume. The free energy cutoff to define the bound state was chosen by convergence of the calculated standard binding free energy (Fig. 9).

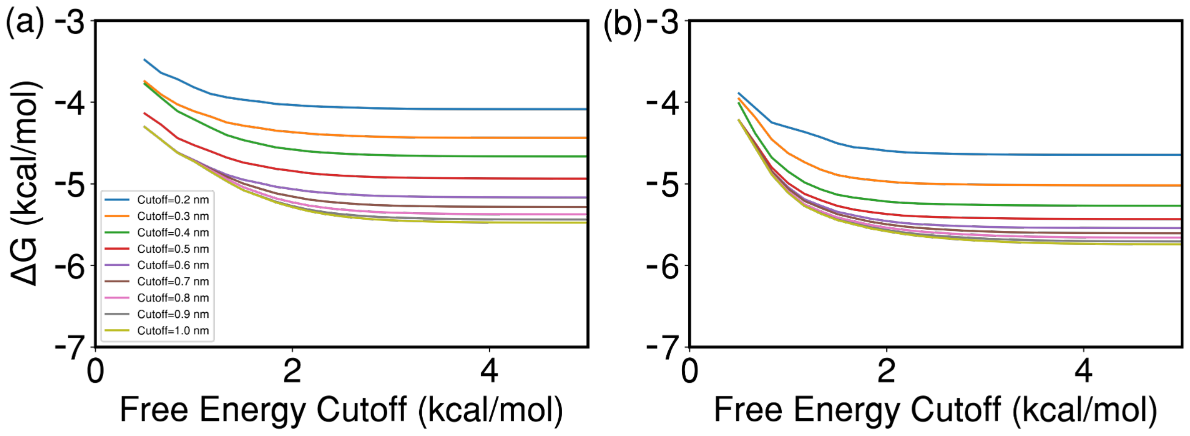


**Figure S9.** Convergence of standard binding free energy with varying free energy and spatial cutoffs for definitions of the bound volume for (a) *At*D14 and (b) *Sh*HTL7.

#### 8 Maximum englobbing regions for binding pocket volume calculation

Binding pocket volume was calculated using the POVME 2.0 software package. The pocket volume calculation first requires the definition of a "maximum englobbing region," which is a field of points spaced in a 1.0 Å grid completely engulfing the binding pocket. The pocket volume is subsequently calculated by removing points within 1.09 Å of any atom and counting remaining points. The ligand was removed for all binding pocket volume calculations.

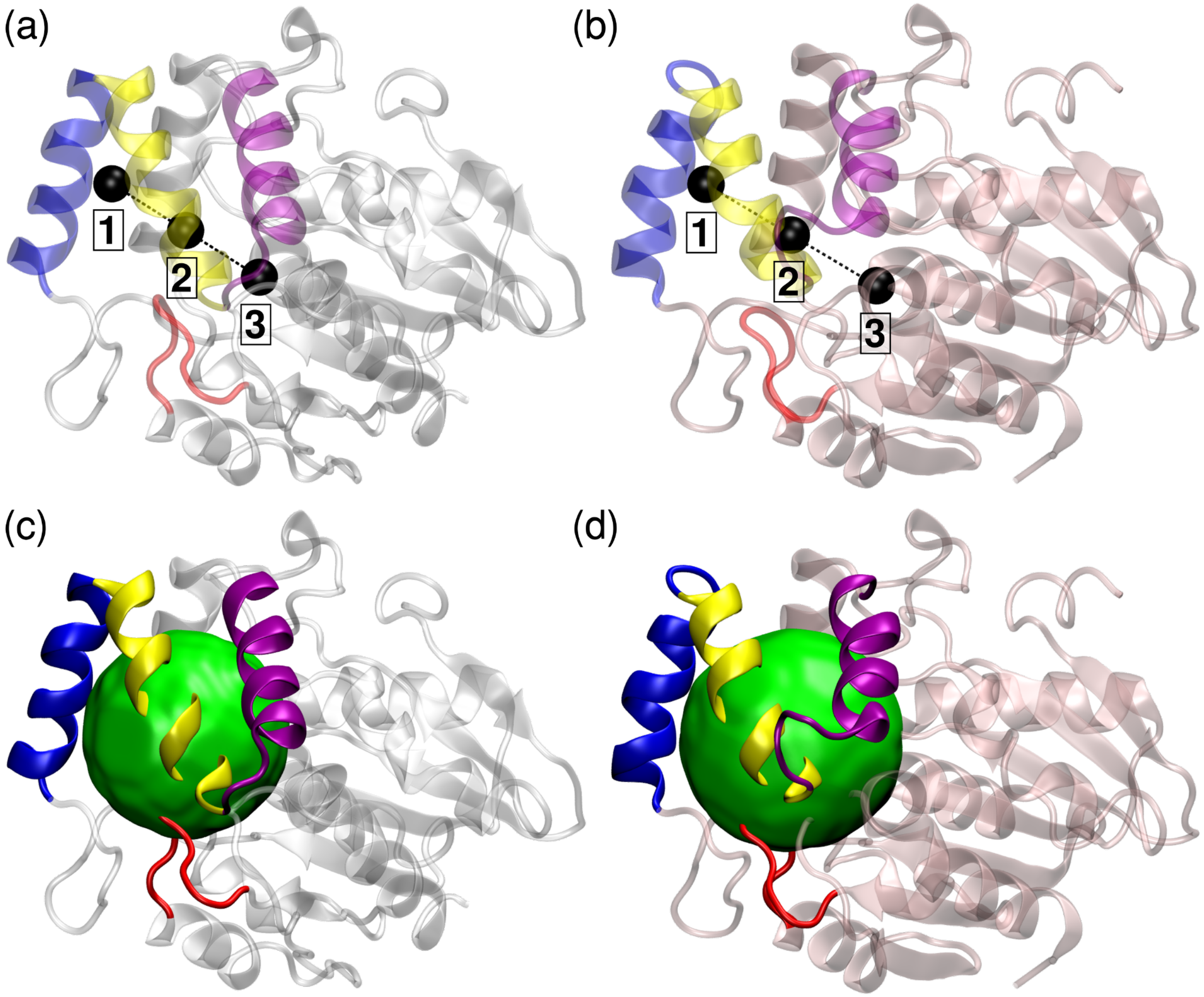


**Figure S10.** Points used to determine maximum englobbing regions for (a) *At*D14 and (b) *Sh*HTL7 pocket volume calculations. Labelled points are (1) the geometric center of the T1 and T2 helices, (2) the center of the inclusion region, and (3) the C-$\alpha$ of S97/95. Green spheres indicate final maximum englobbing regions for (c) *At*D14 and (d) *Sh*HTL7.
